# Multi-organ signaling mobilizes tumor-associated erythroid cells expressing immune checkpoint molecules

**DOI:** 10.1101/2020.07.31.231001

**Authors:** Yasuyo Sano, Toshimi Yoshida, Min-Kyung Choo, Yanek Jiménez-Andrade, Kathryn R. Hill, Katia Georgopoulos, Jin Mo Park

**Affiliations:** Cutaneous Biology Research Center, Massachusetts General Hospital and Harvard Medical School, Charlestown, MA 02129, USA

## Abstract

Hematopoietic-derived cells are integral components of the tumor microenvironment and serve as critical mediators of tumor-host interactions. Cells derived from myeloid and lymphoid lineages perform well-established functions linked to cancer development, progression and response to therapy. It is unclear whether erythroid cells exert such host cell functions in cancer, but emerging evidence points to this possibility. Here we show that tumor-promoting environmental stress and tumor-induced physiological disruption trigger renal erythropoietin production and erythropoietin-dependent expansion of splenic erythroid cell populations in mice. These erythroid cells display molecular features indicative of an immature erythroid phenotype and express immune checkpoint molecules. Erythroid cells with similar properties are present in mouse and human tumor tissues. Antibody-mediated erythropoietin blockade or erythroid cell depletion reduces tumor growth. These findings reveal the potential of erythropoietin and erythroid cells as targets for cancer treatment.

## Introduction

Cells that have acquired cancer-causing genetic alterations are held in check by a multitude of cell-autonomous and non-cell-autonomous mechanisms suppressing neoplastic growth. Additional genetic changes and epigenetic reprogramming allow these cells to evade senescence, programmed death, immune surveillance, and other suppressive mechanisms, thereby permitting their proliferation and progression to a malignant state. Tumor cells interact with host cells throughout the course of their genesis and evolution. Hematopoietic-derived cells are amongst the critical mediators of tumor-host interactions and, depending on the cell type and functional state, exert protumor or antitumor actions in the tumor microenvironment or at remote tissue sites. Knowledge of how host cells of myeloid and lymphoid origin contribute to shaping tumor-associated inflammation and antitumor immunity is rapidly expanding and being translated into cancer prevention and treatment. By contrast, the potential role of host erythroid cells in cancer has been far less explored and remains largely unappreciated. Emerging evidence, however, points to immunomodulatory properties of cells displaying an immature erythroid phenotype in various settings including cancer, lighting up a new path of discovery.

Early evidence suggesting a role for erythroid-lineage cells beyond erythrocyte production came from studies of mechanisms underlying susceptibility to bacterial pathogens. These studies showed that immature erythroid cells were induced upon bacterial infection or accumulated during the neonatal period in mice (1-4), wherein they exerted immunosuppressive actions and impaired antibacterial resistance (2-4). More recently, erythroid cells with similar developmental and functional characteristics were identified in tumor-bearing mice and human cancer patients (5, 6). These erythroid cells were found to promote tumorigenesis by enhancing the survival, migration, and invasive growth of cancer cells (5) and suppressing CD8^+^ T cell-mediated antitumor immunity (6). The discovery of erythroid cells capable of modulating antibacterial and antitumor immune responses indicates that all of the major hematopoietic lineages have evolved to serve immune-related functions. A vast knowledge void still remains, however, regarding how erythroid cells are mobilized in response to pathologic insults and function as regulators of immune responses.

The growth factor erythropoietin (EPO) is essential for erythroid cell development and transduces signals for proliferation and differentiation via the EPO receptor (EPOR) expressed in erythroid progenitors and erythroblasts (7). Recombinant EPO derivatives, known as “erythropoiesis-stimulating agents (ESAs),” have been extensively used to treat anemia resulting from cancer chemotherapy and kidney disease. ESA treatment has proven to be an effective means to manage anemia, yet multiple clinical reports revealed adverse effects among ESA-treated cancer patients such as increased rates of disease progression or recurrence and shortened overall survival (8-10). Consistent with these clinical observations, cell culture and animal model studies detected EPOR expression in and direct EPO actions on cancer cells, proposing a role for EPO signaling in promoting their viability, proliferation, metastatic potential, therapy resistance and stemness (11-16). Evidence against a direct effect of EPO on cancer cells also exists (17, 18), leaving the mechanism for EPO function in cancer still under debate.

In this study, we identify a cascade of signaling events involving multiple organs that link tumor-associated physiological stress to EPO production and EPO-responsive expansion of erythroid cells. Furthermore, we discover that these stress-induced erythroid cells express immune checkpoint molecules, infiltrate tumors, and promote tumor growth.

## Results

### Splenic erythroid cell response to tumor-promoting environmental stress

Ultraviolet-B radiation (UVB) is the major environmental carcinogen for skin cancer and an inducer of skin inflammation. UVB is also known to result in immune dysregulation in ways that cannot be accounted for by its local action in the skin. To investigate systemic effects of skin exposure to UVB, we irradiated shaved back skin of mice with 50 mJ/cm^2^ of UVB, a dose known to elicit acute dermatitis with its local manifestations peaking at 48 to 72 hours after irradiation (19). Included in the analyses for these mice was the detection of bromodeoxyuridine (BrdU) incorporation, a measurement of cell proliferation, in various tissues 96 hours after UVB exposure. This investigation led us to discover massive cell proliferation in the spleen of UVB-irradiated mice but not in the lymph nodes and other locations harboring abundant hematopoietic-derived cells (Fig. 1A). Immunofluorescence analysis of cell type-specific markers revealed that the splenocytes with BrdU incorporation mainly comprised cells expressing the erythroid marker TER119; few of them were stained for CD3, B220, or CD11b (markers for T cells, B cells, and myeloid cells, respectively; Fig. 1B). UVB-induced splenic erythroid cell proliferation was confirmed by click chemistry-based detection of 5-ethynyl-2′-deoxyuridine (EdU) incorporation in splenocytes expressing not only TER119 but also CD71, a marker associated with earlier stages of erythroid cell differentiation (Supplemental Fig. 1).

**Figure 1.**
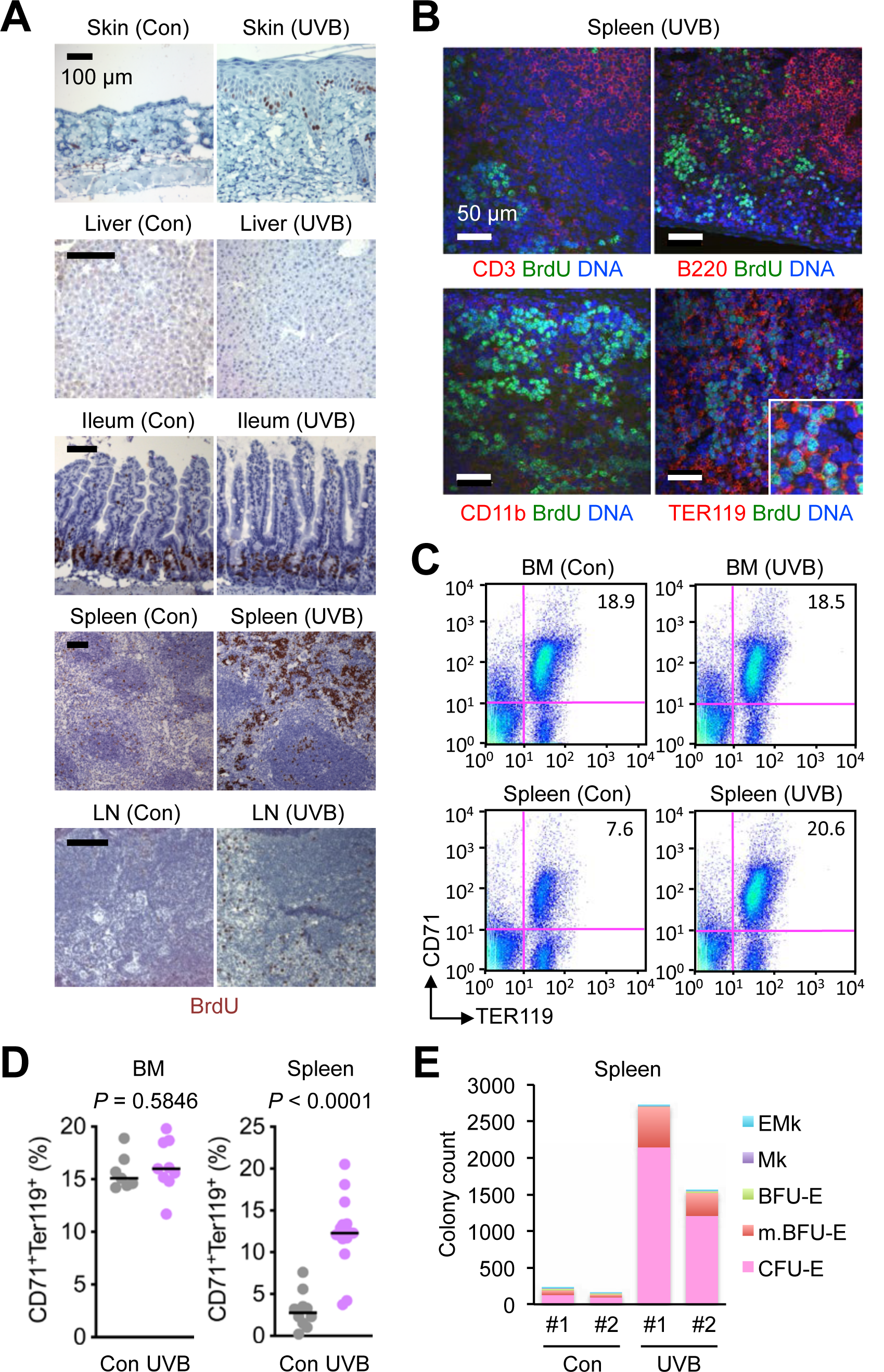
Skin exposure to UVB results in expansion of splenic erythroid cell populations in mice. (**A** and **B**) Tissues were isolated from C57BL/6 mice 4 days after their shaved back skin was exposed to UVB (50 mJ/cm^2^). BrdU was injected intraperitoneally 3 hours before tissue isolation. Tissue sections were analyzed by immunohistochemistry (**A**) and immunofluorescence along with DNA counterstaining (**B**). Tissues from control (Con) mice (shaved, unirradiated, and BrdU-injected) were prepared and analyzed in parallel. LN, lymph node. Data are representative of three experiments. (**C** and **D**) Bone marrow (BM) cells and splenocytes were prepared from unirradiated and UVB-irradiated mice as in **A** and analyzed by antibody staining and flow cytometry (**C**). CD71^+^TER119^+^ cell percentages in individual mice (circle) and their medians (line) are shown (**D**). Data are representative of two experiments. (**E**) Splenic nucleated cells prepared from mice left unirradiated and irradiated with UVB as in **A** were plated for the formation of erythroid-megakaryocyte (EMk), megakaryocyte (Mk), burst-forming unit-erythroid (BFU-E), mature BFU-E (m.BFU-E) and colony-forming unit-erythroid (CFU-E) colonies. Colony counts indicate numbers of colonies per 5 × 10^5^ plated cells for individual mice (#1 and #2) in each group. Data are representative of two experiments.

Skin exposure to UVB triggered expansion of erythroid cell populations in the spleen but not the bone marrow (Fig. 1, C and D). The expanded erythroid cell populations expressed both CD71 and TER119, suggesting an immature, erythroid progenitor- or erythroblast-like state. In line with this finding, we observed that the spleen of UVB-irradiated mice contained greater numbers of cells with the potential for erythroid colony formation (Fig. 1E).

### Expression of immune checkpoint molecules in stress-induced splenic erythroid cells

We performed RNA sequencing (RNA-Seq) analysis of UVB-induced splenic erythroid cells versus bone marrow erythroid cells engaged in steady-state hematopoiesis to identify differences in their phenotype and functional capabilities (Fig. 2A). This gene expression profiling showed high expression of lineage-specific genes, such as *Tfrc, Gata1*, and *Klf1*, in both groups of erythroid cells. In addition, comparison of the two transcriptomes revealed genes uniquely expressed in UVB-induced splenic erythroid cells. Gene ontology enrichment analysis associated these differentially expressed genes with immunomodulatory properties as their functional attributes (Fig. 2B). Most notable among the genes with such functional annotations were those encoding immune checkpoint molecules and other similar regulatory proteins, including *Cd274* (encoding PDL1) and *Cd244* (encoding 2B4). Flow cytometry analysis confirmed preferential expression of PDL1 and 2B4 proteins in UVB-induced splenic CD71^+^TER119^+^ cells relative to their counterparts in the steady-state bone marrow (Fig. 2, C and D). These findings indicated that stress-induced immature erythroid cells might not merely serve as precursors of erythrocytes but perform specialized regulatory functions in immunity.

**Figure 2.**
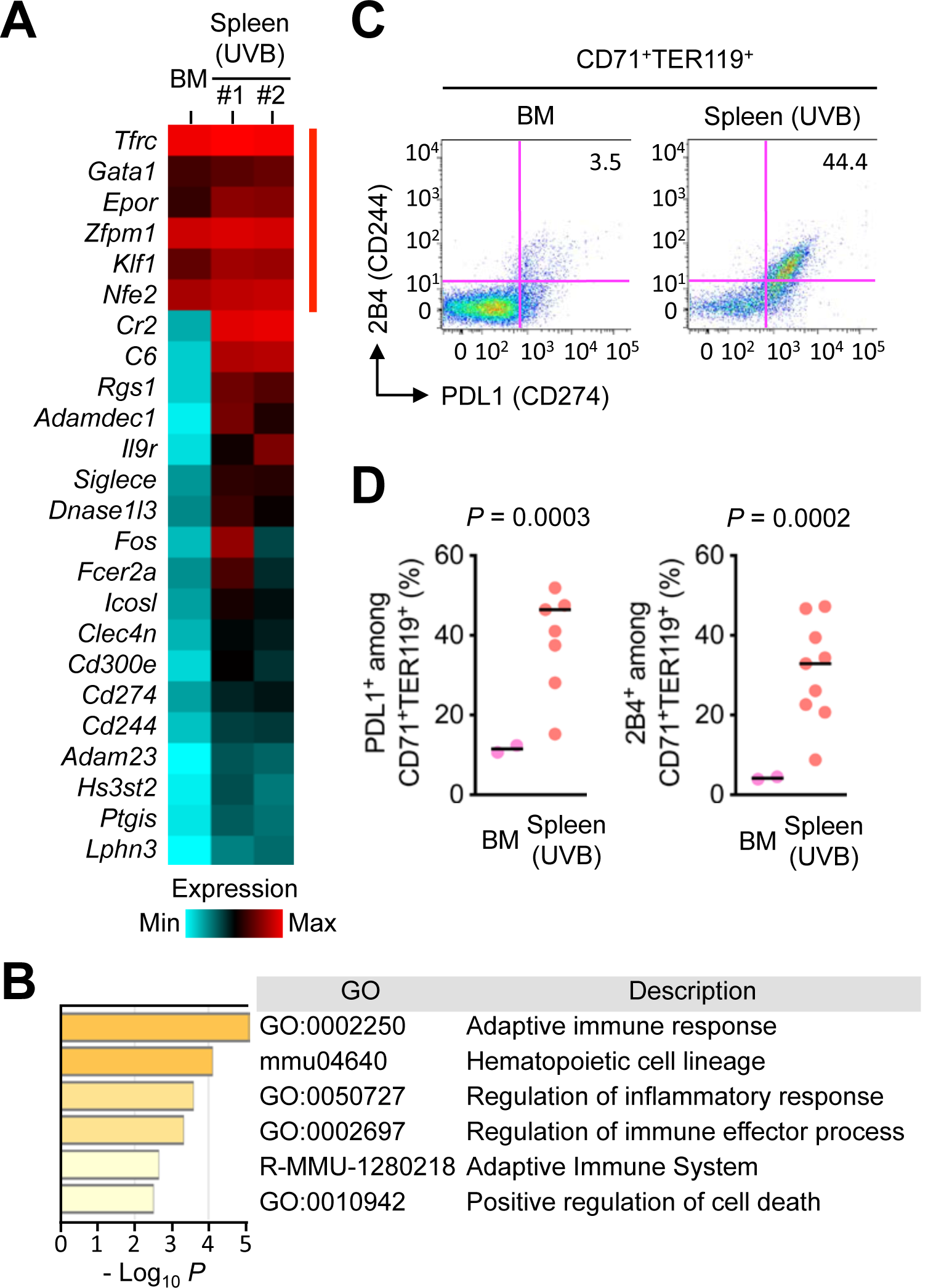
UVB-induced splenic erythroid cells express immune checkpoint molecules. (**A** and **B**) CD71^+^TER119^+^ cells were purified from the spleen of mice irradiated with UVB as in **Fig. 1A** and subjected to transcriptome analysis by RNA-Seq (**A**). CD71^+^TER119^+^ cells purified from the bone marrow (BM) of unirradiated mice were analyzed in parallel. Relative mRNA amounts for erythroid signature genes (indicated by red line on the right) and genes differentially expressed in the two erythroid cell populations are presented in color-coded arbitrary units. Functional attributes represented by the differentially expressed genes (corresponding to *Cr2* through *Lphn3* in the gene list on the left in **A**) were ranked using Metascape, a tool for gene ontology enrichment analysis (**B**). The sources of GO terms used in Metascape analysis included functional categories in the Gene Ontology Resource, KEGG Pathway and Reactome Pathway databases. *P* values indicate the statistical significance of the GO terms being enriched in the input list. (**C** and **D**) BM cells and splenocytes prepared as in **A** were analyzed by antibody staining and flow cytometry (**C**). The percentages of PDL1^+^ and 2B4^+^ cells among CD71^+^TER119^+^ cells in individual animals (circle) and their medians (line) are shown (**D**). Data are representative of two experiments.

### Molecular pathway linking tumor-promoting environmental stress to splenic erythroid cell proliferation

The discovery of the erythroid response to UVB exposure directed our attention to EPO. In adult mammals, EPO is produced mainly in the kidney in response to reduced oxygen transport and acts on hematopoietic tissues as a circulating cytokine. We found that UVB-irradiated mice produced increased amounts of circulating EPO (Fig. 3A). Correspondingly, these mice exhibited robust induction of EPO synthesis in renal peritubular cells (Fig. 3B). Antibody-mediated EPO neutralization suppressed UVB-induced expansion of nucleated TER119^+^ cell clusters in spleen sections (Fig. 3, C and D), demonstrating an indispensable role for EPO in UVB-responsive splenic erythroid cell proliferation.

**Figure 3.**
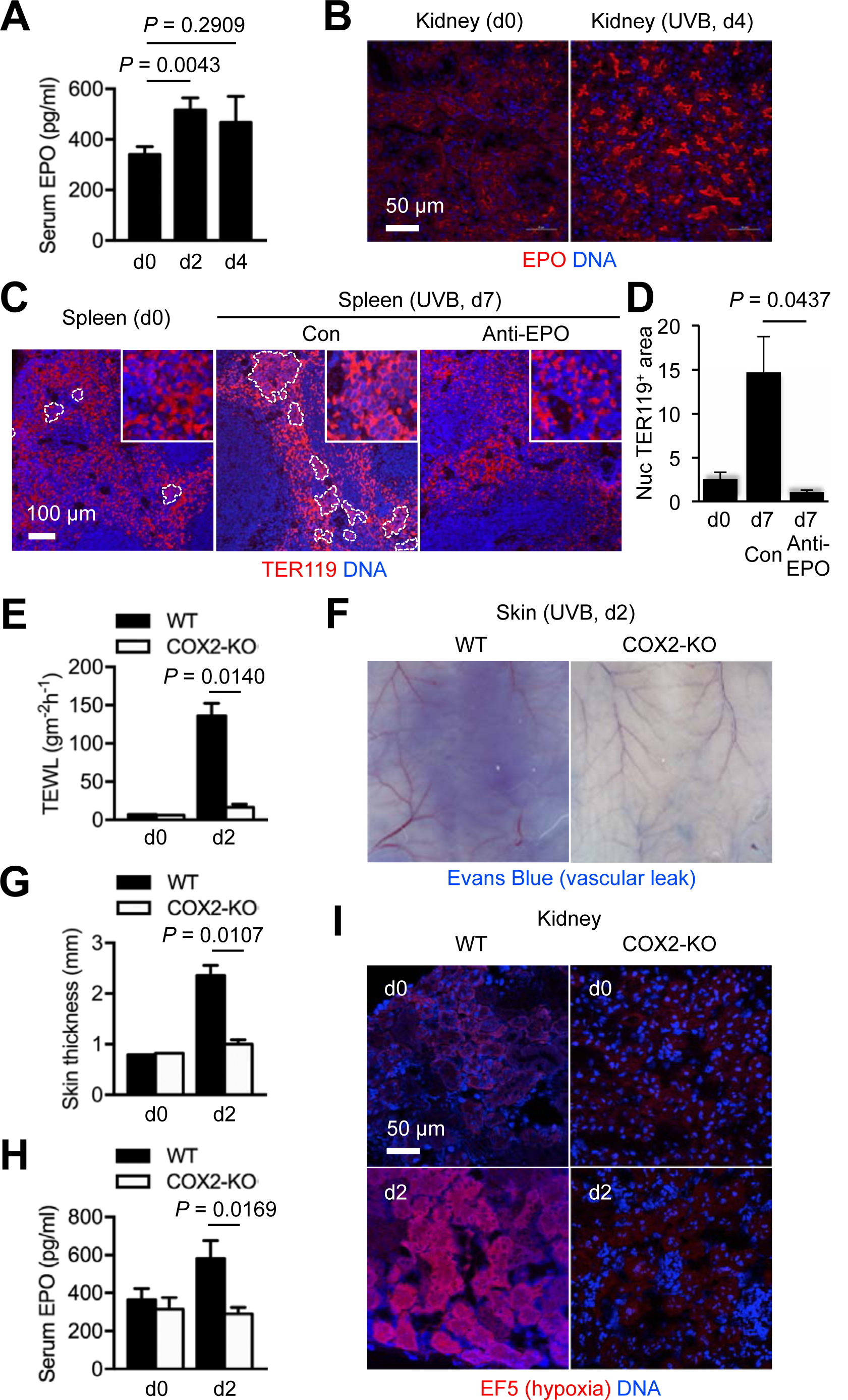
UVB-responsive splenic erythroid expansion in mice depends on inflammation-induced renal EPO production. Mice were left unirradiated or irradiated with UVB as in **Fig. 1A**. Peripheral blood and tissues were prepared before and at the indicated time points after irradiation (dn, day n) for further analysis. (**A**) Serum EPO concentrations in C57BL/6 mice (*n* = 16, 19, and 5 for d0, d2, and d4, respectively) were determined by ELISA. Data are representative of five experiments. (**B**-**D**) Tissue sections from C57BL/6 mice were analyzed by immunofluorescence and DNA counterstaining 4 days (**B**) and 7 days (**C** and **D**) after UVB exposure. An anti-EPO antibody and isotype-matched control (Con) immunoglobulin were administered to mice on d1, d3, and d5 after UVB irradiation (**C** and **D**). Clusters of nucleated TER119^+^ cells are demarcated by dotted line (**C**). Areas of these erythroblast-like cell clusters relative to total TER119^+^ areas in image fields (*n* = 4) were quantified (**D**). Nuc, nucleated. Data are representative of three (**B**) and two (**C** and **D**) experiments. (**E**-**I**) UVB-induced changes in transepidermal water loss (TEWL; **E**), vascular diameter and permeability (**F**), skin thickness (**G**), serum EPO concentration (**H**), and renal hypoxia (**I**) in WT and COX2-KO mice were compared. Evans Blue (**F**) and EF5 (**I**) were injected into mice 30 minutes before tissue isolation. The inner side of back skin flaps was photographed (**F**). EF5 adduct formation in hypoxic tissue areas was detected by immunofluorescence with an EF5-specific antibody (**I**). Data are representative of two experiments.

We suspected inflammation as a possible link between skin exposure to UVB and renal EPO induction. The dose of UVB used in this study has been shown to induce epidermal expression of cyclooxygenase-2 (COX2), an enzyme catalyzing prostaglandin production, and bring about COX2-dependent acute dermatitis in mice (19). Using COX2-knockout (KO) mice, we examined whether preventing dermatitis in UVB-irradiated mice would attenuate EPO production. Tissue changes linked to UVB-induced inflammation, including epidermal barrier disruption (elevated transepidermal water loss), vasodilation, and vascular permeability increase (edema formation), were substantially reduced in COX2-KO skin (Fig. 3, E-G). Importantly, UVB irradiation failed to induce EPO production in the absence of these inflammatory responses (Fig. 3H). UVB-induced skin inflammation likely led to a decrease in total plasma volume and an increase in blood flow rate in the skin due to vascular leakage and vasodilation, respectively. These two hemodynamic changes together were expected to result in reduced rates of blood flow in and oxygen supply to the kidney, leaving renal tissue in microhypoxia, a condition known to trigger EPO gene expression (20). To test this possibility, we administered EF5, a nitroimidazole compound forming chemical adducts in hypoxic tissues (21), to UVB-irradiated mice and performing EF5-specific immunofluorescence analysis of their kidney sections (Fig. 3I). Renal hypoxia peaked at 48 hours after UVB exposure, but this response was absent in COX2-KO mice (Fig. 3I). Taken together, these findings highlighted the involvement of multi-organ signaling events in stress-induced expansion of splenic erythroid cells.

### Splenic erythroid cell response to tumor growth in remote locations

Established tumors create a tissue environment with a chronic inflammatory tone and activated vasculature, likely causing hemodynamic changes and renal EPO induction similarly to UVB-exposed skin. Mice bearing tumors formed from mouse melanoma B16 cells and mouse colon adenocarcinoma MC38 indeed had increased serum EPO concentrations and renal EPO immunofluorescence signals compared to control samples (Fig. 4, A and B). B16 and MC38 tumor-bearing mice also exhibited nucleated TER119^+^ cells with BrdU incorporation in the spleen (Supplemental Fig. 2A) and expanded CD71^+^TER119^+^ splenocyte populations (Fig. 4, C and D; Supplemental Fig. 2B). These tumor-induced erythroid cells appeared to have close parallels with other recently reported erythroid-lineage cells that were associated with hepatocellular carcinoma, lung carcinoma, and other cancer types and found to possess immunosuppressive and tumor-promoting capacities (5, 6). To verify the ability of erythroid cells to promote tumor growth, we examine the effect of antibody-mediated EPO neutralization and erythroid cell depletion on B16 and MC38 tumor growth. Anti-EPO and anti-CD71 antibodies administered to tumor-bearing mice were capable of preventing erythroid expansion and depleting expanded erythroid populations, respectively (Supplemental Fig. 3). Treatment with these antibodies produced moderate antitumor effects, slowing subcutaneous growth of B16 and MC38 tumors (Fig. 4, E and F).

**Figure 4.**
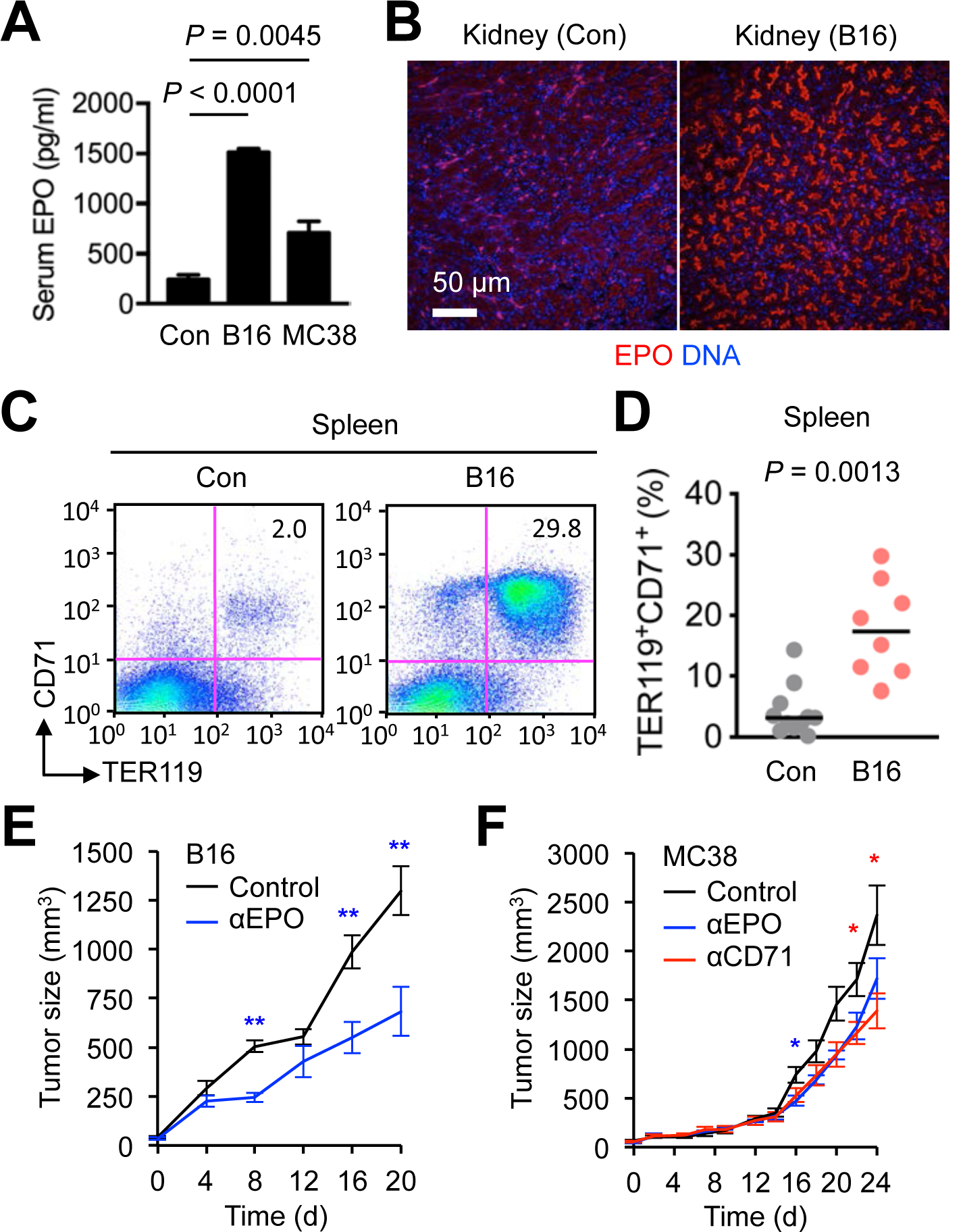
Tumor growth results in expansion of splenic erythroid cell populations in mice. (**A**-**D**) Peripheral blood and tissues were prepared from mice bearing subcutaneous B16 and MC38 tumors of 1,000-2,500 mm^3^ in size and control (Con) mice without tumor growth. Serum EPO concentrations (*n* = 5, 3, and 7 for Con, B16 and MC38 mice, respectively) were determined by ELISA (**A**). Kidney sections were analyzed by immunofluorescence and DNA counterstaining (**B**). Splenocytes were analyzed by antibody staining and flow cytometry (**C**). CD71^+^TER119^+^ cell percentages in individual mice (circle) and their medians (line) are shown (**D**). Data are representative of three experiments. (**E** and **F**) B16 and MC38 cells were injected into C57BL/6 mice (*n* = 6-8) to form subcutaneous tumors. Anti-EPO and anti-CD71 antibodies and isotype-matched control (Con) immunoglobulin were administered to mice (every other day, three times in total, and the first dose given when the size of B16 and MC38 tumors reached 100 mm^3^ and 300 mm^3^, respectively). d, day. **, *P* < 0.01; *, *P* < 0.05. Data are representative of three (**E**) and two (**F**) experiments.

### Erythroid cell infiltration of tumor tissue

Given the reports of non-erythropoietic EPO functions and non-erythroid EPOR expression, we explored the possibility that EPO promoted tumor growth through acting on EPOR ectopically expressed in cancer cells apart from its ability to mobilize erythroid cells. We first examined EPOR expression in tumor sections by immunofluorescence analysis. EPOR expression was detectable in B16 tumors growing in mice but restricted to a distinct fraction of cells scattered across the intratumoral space (Fig. 5A, left). Human melanoma Hs944T tumors growing in immunodeficient host mice (*Rag2*^*-/-*^*Il2rg*^*-/-*^) exhibited a similar EPOR expression pattern with EPOR^+^ cells detected only in a subset of cells in tumor sections (Fig. 5A, right). Almost all EPOR^+^ cells in tumors were nucleated and stained for TER119, whereas EPOR expression was undetectable among anucleated TER119^+^ cells (Fig. 5, B and C). Flow cytometry analysis revealed that B16 tumors contained cells expressing CD71 and TER119, either alone or together, with EPOR expression highest in the CD71^+^TER119^+^ double-positive population (Fig. 5D). CD71^+^TER119^+^ cells purified from tumors by flow sorting had an intact nucleus and displayed a morphology similar to that of basophilic erythroblasts (Fig. 5, E and F). Infiltration of CD71^+^ cells and cells expressing CD235A, another human erythroid marker, was also detected in clinical tumor samples from patients with various cancer types (Fig. 5G; Supplemental Fig. 4).

**Figure 5.**
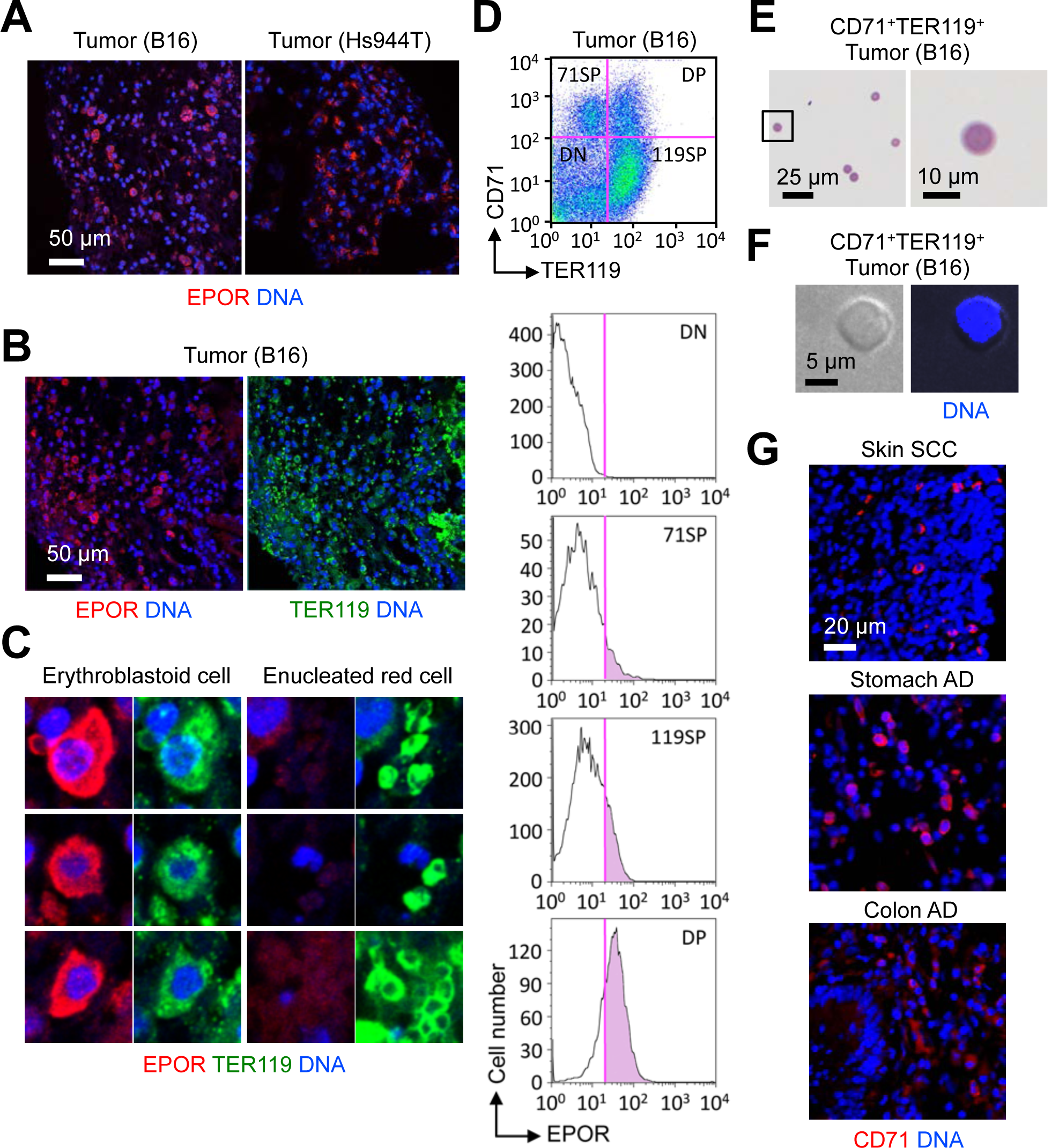
EPOR expression in tumor tissues is restricted to tumor-associated erythroid cells. (**A**-**C**) Sections of B16 and Hs944T tumors growing in WT and RAG2/γ_c_-double KO mice, respectively, were analyzed by immunofluorescence and DNA counterstaining. Data are representative of two experiments. (**D**) Total live cells from a B16 tumor were analyzed by antibody staining and flow cytometry. Data are representative of three experiments. (**E** and **F**) CD71^+^TER119^+^ cells were purified from B16 tumors and analyzed by May-Giemsa staining (**E**) and DNA fluorescence (**F**). Data are representative of two experiments. (**G**) Human tumor sections were analyzed by immunofluorescence and DNA counterstaining.

### Expression of immune checkpoint molecules in tumor-associated erythroid cells

To obtain information on the phenotype of tumor-induced erythroid cells and discern their functional potential related to cancer, we performed RNA-Seq analysis of erythroid cells purified from the spleen of B16 tumor-bearing mice and compared their transcriptome with that of bone marrow erythroid cells from control mice without tumors (Fig. 6A). Tumor-induced erythroid cells were found to express erythroid signature genes as well as a distinct subset of genes that set them apart from erythroid cells in the steady-state bone marrow. Similar to erythroid cells purified from UVB-irradiated mice, tumor-induced erythroid cells exhibited high expression of *Cd274* and other genes encoding immune checkpoint molecules. There were a group of genes highly expressed in tumor-induced but not UVB-induced erythroid cells, such as *Fam132b* (also known as *Erfe*; encoding erythroferrone) and *Osm* (encoding oncostatin-M). These gene expression profiles indicated that erythroid cell populations induced by skin exposure to UVB and by tumor growth might share immunomodulatory properties while possessing distinct functional capabilities conferred by the genes uniquely expressed in either cell population.

**Figure 6.**
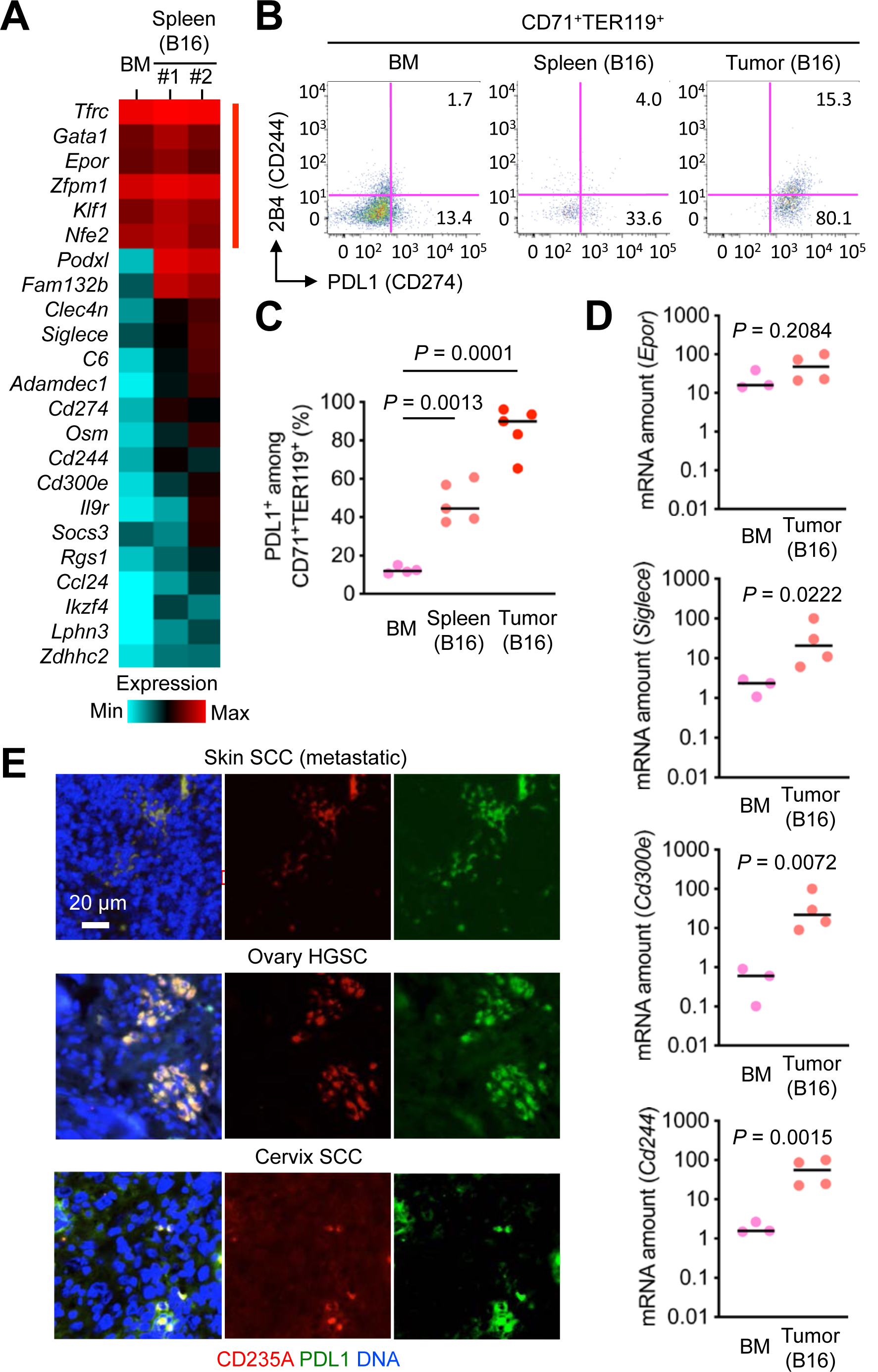
Tumor-associated erythroid cells express immune checkpoint molecules. (**A**) CD71^+^TER119^+^ cells were purified from the spleen of B16 tumor-bearing mice and subjected to transcriptome analysis by RNA-Seq. CD71^+^TER119^+^ cells purified from the bone marrow (BM) of mice without tumor growth were analyzed in parallel. Relative mRNA amounts for erythroid signature genes (indicated by red line on the right) and genes differentially expressed in the two erythroid cell populations are presented in color-coded arbitrary units. (**B** and **C**) BM cells and splenocytes prepared as in **A** and total cells from B16 tumors were analyzed by antibody staining and flow cytometry (**B**). The percentages of PDL1^+^ cells among CD71^+^TER119^+^ cells in individual animals (circle) and their medians (line) are shown (**C**). Data are representative of two experiments. (**D**). CD71^+^TER119^+^ cells were purified from the BM of mice without tumor growth and B16 tumors growing in mice and subjected to qPCR analysis. Data were obtained from one experiment. (**E**). Human tumor sections were analyzed by immunofluorescence and DNA counterstaining.

We examined whether erythroid cells isolated from B16 tumors also expressed genes encoding immune checkpoint molecules and other genes specifically associated with tumor-induced splenic erythroid cells. Flow cytometry analysis showed that the *Cd274* product PDL1 was highly expressed on tumor-infiltrating CD71^+^TER119^+^ cells (Fig. 6, B and C). Quantitative PCR analysis of CD71^+^TER119^+^ cells purified from B16 tumors by flow sorting revealed greater amounts of transcripts of *Siglec3, Cd300e*, and *Cd244*, whereas *Epor* expression in these cells was comparable to that in control erythroid cells from the bone marrow (Fig. 6D). Cells expressing the erythroid marker CD235A together with PDL1 were detected in human tumor samples from cancer patients (Fig. 6E). Our findings suggest tumor-associated erythroid cells and immunomodulatory proteins expressed in these cells as novel mediators of tumor-host interactions and actionable targets for cancer therapy.

## Discussion

We have shown that erythroid cells serving immunomodulatory and tumor-promoting functions are mobilized in response to UVB exposure, a form of tumor-promoting environmental stress, and tumor-induced physiological disruption. Inflammation-driven hemodynamic changes appear to play a key role in inducing renal EPO production and EPO-dependent expansion of these erythroid cell populations. Our findings represent novel mechanisms by which preneoplastic tissues and established tumors communicate with and leverage support from other tissues in remote locations. The multi-organ signaling events shown to occur in carcinogen-exposed or tumor-bearing mice culminate in the induction of erythroid cells expressing unique immunomodulatory genes including those encoding immune checkpoint molecules.

Previous findings showing unintended clinical effects of ESAs and EPOR expression in non-erythroid cells hinted at EPO functions beyond erythropoiesis. Some of these non-erythropoietic functions appeared to underlie clinically beneficial effects, such as those related to tissue protection and attenuated inflammatory responses (22, 23). Animal model studies showed that EPO protected neurons, cardiomyocytes, pancreatic β cells, and other non-erythroid cells against injuries of various nature (24-27), and alleviated ischemic cerebral inflammation and colitis (28, 29). On the other hand, the majority of the non-erythropoietic EPO functions in cancer were linked to adverse effects of ESAs in cancer patients (8-10). These effects were thought to derive from ESA-induced signaling via cancer cell-expressed EPOR. Based on the results of this study, we offer an alternative explanation, proposing EPO-responsive immunomodulatory erythroid cells as a mediator of the protumor effects of endogenously produced EPO or clinically administered ESAs. We do not rule out, however, the contribution of EPO signaling via EPOR expressed in cancer cells and other tumor-associated host cell types.

Many of the cancer immunotherapies in use or development achieve their efficacy by reinvigorating cytotoxic lymphocytes in a dysfunctional state and restoring their ability to detect and destroy cancer. Other approaches of cancer immunotherapy are aimed at eliminating or inactivating immunosuppressive myeloid cells or augmenting phagocytic cancer cell clearance. Findings from this study will prompt a new line of investigation for assessing host erythroid cells as a target for cancer therapy. Our findings on the signaling mechanisms linking tumor growth to erythroid cell mobilization and the phenotypic differences between tumor-associated erythroid cells and bone marrow erythroid cells engaged in hematopoiesis will help devise new therapeutic strategies. The transcriptomes of UVB- and tumor-induced erythroid cells were found to contain transcripts suggestive of their immunomodulatory properties. In addition to PDL1, a major immune checkpoint molecule targeted by therapeutics in clinical use (30-33), several of the identified genes encode proteins that have been shown to control antitumor immunity in preclinical studies (e.g. 2B4, SIGLEC-E, interleukin-9 receptor; 34-36) and molecular markers whose expression in the tumor microenvironment is prognostic for a poor clinical outcome (e.g. PODXL, ADAMDEC1; 37, 38). The knowledge of their expression in erythroid cells provides the context in which they serve cancer-related functions and enriches our understanding of the tumor immune microenvironment.

## Materials and Methods

### Mice

C57BL/6 mice (Jackson Laboratory), COX2-KO mice (Taconic Biosciences) in a mixed B6;129P2 background, and RAG2/γ_c_-double KO mice in a C57BL/6 background (Taconic Biosciences) were maintained and used in experiments in a specific pathogen-free condition. All animal experiments were conducted under an Institutional Animal Care and Use Committee-approved protocol.

### UVB irradiation of mouse skin and skin analysis

Shaved and depilated back skin of 2–3-month-old mice was irradiated with 50 mJ/cm^2^ of UVB using UVB bulbs (Southern NE Ultraviolet) and a Kodacel filter (Eastman Kodak). UVB dose was monitored with a radiometer (International Light). BrdU (50 mg/kg; Sigma) and EdU (50 mg/kg; Thermo Fisher Scientific) were administered to mice intravenously 3 hours before euthanasia and tissue isolation. Evans Blue (0.1 ml of 1% solution in 0.9% saline per animal; Sigma) and EF5 (0.2 ml of 10 mM solution in 0.9% saline per animal; a gift from Cameron Koch, University of Pennsylvania) were administered to mice intravenously 30 minutes before euthanasia and tissue isolation. The rate of transepidermal water loss was measured with a vapometer (Delfin Technologies). The thickness of lifted back skin was measured with a caliper (Mitutoyo).

### Tumor formation in mice and tumor growth measurement

B16, MC38, and Hs944T cells (1 × 10^6^ cells per animal) were injected into the right hind legs of 8-week-old mice to establish tumors. Tumor size was manually measured with a caliper.

### EPO neutralization and erythroid cell depletion in vivo

Anti-EPO (148438, R&D Systems) and anti-CD71 (8D3, BioRad Laboratories) antibodies and isotype-matched control immunoglobulin (0.25 mg per animal) were administered to mice every other day and three times in total. The first dose was given 24 hours after UVB exposure and when tumors grew to 100 mm^3^ (B16) and 300 mm^3^ (MC38) in size.

### Histology and immunofluorescence analysis

Mouse organ and tumor tissues were fixed with formalin and embedded in paraffin. Tissue sections were analyzed by hematoxylin and eosin staining or by immunofluorescence analysis using primary antibodies in conjunction with Alexa Fluor 488/594-conjugated secondary antibodies (Thermo Fisher Scientific) and Hoechst 33342 (Sigma). Primary antibodies specific to the following antigens were used in immunofluorescence analysis after 1:50 to 1:1000 dilution: BrdU, CD3e (SP7; Abcam), B220 (RA3-6B2; BD Biosciences), CD11b (M1/70; eBioscience), TER119 (TER-119; BD Biosciences), EPO (Santa Cruz Biotechnology), EPOR (Abcam), human CD71 (Abcam), human CD235A (Abcam), human PDL1 (E1L3N; Cell Signaling Technology), and EF5 (ELK3-51; a gift from Cameron Koch, University of Pennsylvania). BrdU in tissue sections were analyzed using the BrdU In Situ Detection Kit (BD Biosciences). De-identified human tumor sections for immunofluorescence analysis were obtained from commercial sources (US Biomax). Immunohistochemistry images of human tumor sections were obtained from the Human Protein Atlas project (39).

### Flow Cytometry and cell sorting

Single-cell suspensions obtained from mouse tissues were incubated with Fc receptor-blocking anti-CD16/CD32, stained with fluorescent-conjugated antibodies, and analyzed by flow cytometry using FACSCanto (BD Biosciences) and FlowJo software (Tree Star) or subjected to cell sorting using SH800 (Sony). Antibodies specific to the following antigens were used in flow cytometry: CD71 (C2/C2F2; BD Biosciences), TER119 (TER-119; BD Biosciences), PDL1 (10F.9G2; BioLegend), and 2B4 (eBio244F4; eBioscience).

### Methylcellulose colony forming cell assay

5 × 10^5^ nucleated splenocytes were cultured with Methocult M3434 medium (STEMCELL Technologies) supplemented with human thrombopoietin (50 ng/ml), human interleukin-11 (50 ng/ml), and mouse granulocyte-macrophage colony-stimulating factor (10 ng/ml) in 35-mm culture plates. Colony-forming unit-erythroid (CFU-E) and mature burst-forming unit-erythroid (m.BFU-E) colonies were scored after 2-3 days of culture according to the manufacturer’s instructions. The other types of colonies were scored after 5-14 days of culture according to previously described criteria (40) and confirmed by May-Giemsa staining (Harleco) of cytospins of individual colonies.

### RNA analysis

Total RNA was isolated using the Trizol Reagent (Thermo Fisher Scientific) and subjected to cDNA synthesis using the SuperScript IV VILO Master Mix (Thermo Fisher Scientific). Relative transcript abundance was determined by real-time quantitative PCR using the SYBR Green PCR Master Mix (Applied Biosystems) and gene-specific primers. For transcriptome analysis, total RNA isolated with the Trizol Reagent was purified using the RNeasy Mini Kit (Qiagen) and subjected to sequencing library construction (TruSeq Stranded mRNA Prep, Illumina) and 50-cycle single-end sequencing (HiSeq, Illumina) at the MGH Sequencing core facility. Sequencing reads were mapped to a Mus musculus reference genome (assembly mm10). Gene expression counts were calculated using the program HTSeq (41). Gene sets were functionally annotated using the Metascape tool (42). All RNA-Seq data are available in the NCBI GEO database (GSE113874).

### Statistical Analysis

Data values are expressed as mean ± S.E.M. unless indicated otherwise. *P* values were obtained with the unpaired two-tailed Student’s *t*-test.

## Acknowledgments

We thank Cameron Koch for materials. This study was supported by the National Institutes of Health grant CA182405 (JMP).

