## Supplementary material for "Multi-organ signaling mobilizes tumor-associated erythroid cells expressing immune checkpoint molecules": Figures S1-S4

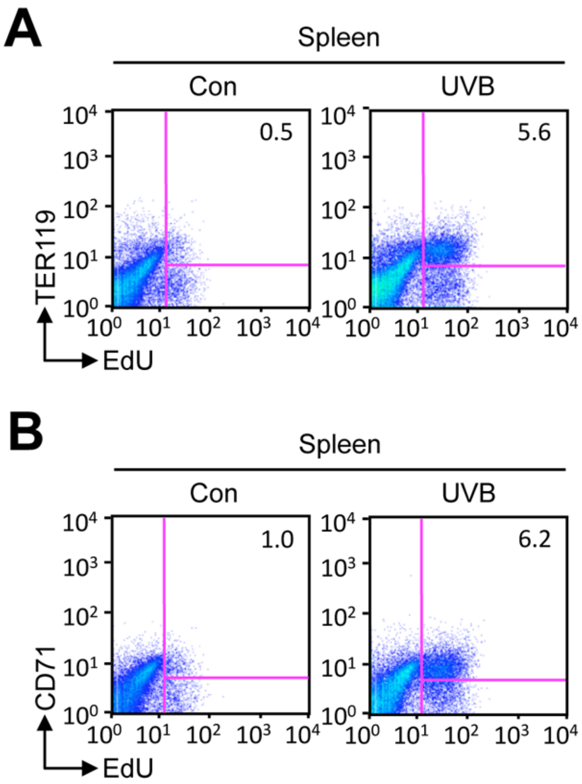


**Figure S1. Skin exposure to UVB induces splenic erythroid cell proliferation in mice.**

(**A** and **B**) Splenocytes prepared from C57BL/6 mice 4 days after their shaved back skin was exposed to UVB (50 mJ/cm$^2$) were analyzed by antibody staining and flow cytometry. EdU was injected intraperitoneally 3 hours before splenocyte isolation. Splenocytes from control (Con) mice (shaved, unirradiated, and EdU-injected) were prepared and analyzed in parallel. Data are representative of two experiments.

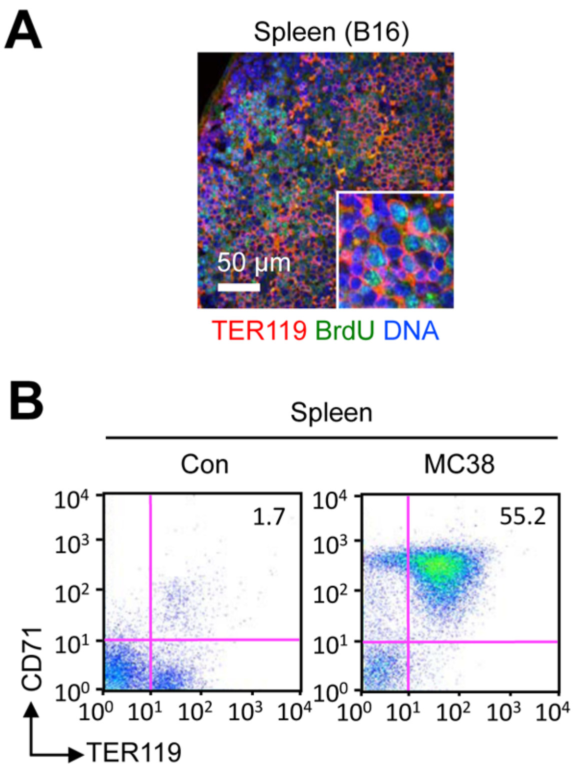


**Figure S2. Tumor growth induces splenic erythroid cell proliferation in mice.**

(**A**) The spleen of a mouse bearing subcutaneous B16 tumors of approximately 2,000 $mm^3$ in size was isolated 3 hours after intraperitoneal BrdU injection. A spleen section was analyzed by immunofluorescence along with DNA counterstaining. Data are representative of two experiments.

(**B**) Splenocytes prepared from a mouse bearing subcutaneous MC38 tumors of 2,000-2,500 $mm^3$ in size and a control (Con) mouse without tumor growth were analyzed by antibody staining and flow cytometry. Data are representative of three experiments.

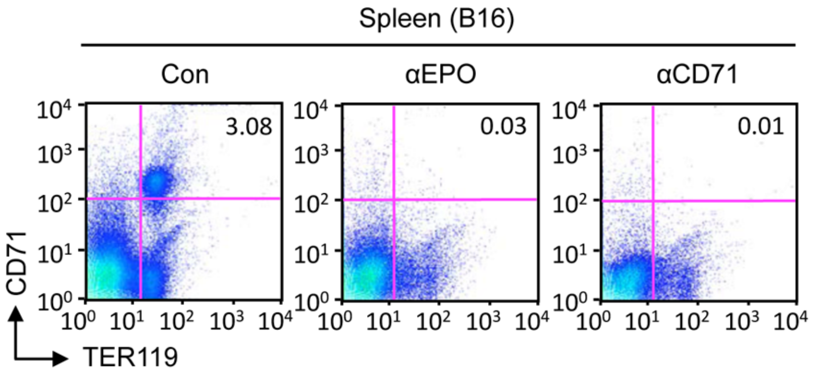


**Figure S3. Anti-EPO and anti-CD71 antibodies prevent the induction of or deplete splenic erythroid cells in tumor-bearing mice.**

Anti-EPO and anti-CD71 antibodies and isotype-matched control (Con) immunoglobulin were administered to B16 tumor-bearing mice every other day and three times in total with the first dose given when tumors grew to 100 $mm^3$ in size. Splenocytes prepared from these mice were analyzed by antibody staining and flow cytometry. Data are representative of three experiments.

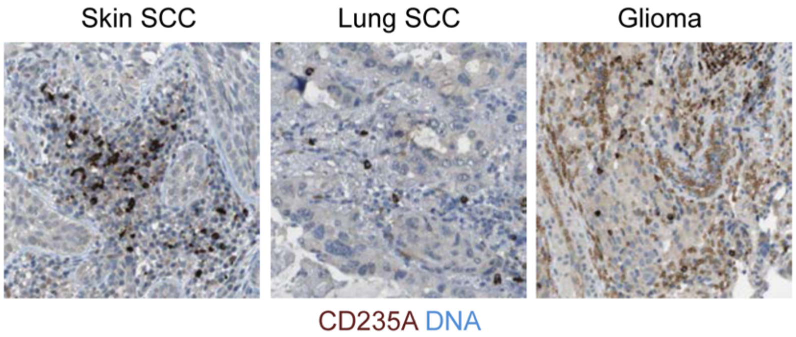


**Figure S4. CD235A immunostaining identifies tumor-associated erythroid cells in human cancer tissues.**

Immunohistochemistry images from the Human Protein Atlas project reveal CD235A-expressing cells in human tumors.
